# Mutations in apicoplast rRNA genes are associated with clindamycin resistance and impair the ability of malaria parasites to infect mosquitoes

**DOI:** 10.64898/2026.04.07.716898

**Authors:** Jessica L. Home, Lee Ming Yeoh, Geoffrey I. McFadden, Christopher D. Goodman

## Abstract

Drug resistance hampers malaria treatment and control. Resistance to nearly all clinically used antimalarials has emerged and spread globally. With multi-drug-resistant parasites now on the rise, understanding resistance mechanisms, and their ability to spread, is crucial for optimal treatment and control strategies. Clindamycin is an apicoplast-targeting antimalarial used as a partner compound in second-line treatment combinations, but mechanisms of clindamycin resistance remain largely unexplored, and it is unclear whether resistant parasites could spread readily. We selected *in vitro* for clindamycin resistance in African and Southeast Asian strains of *Plasmodium falciparum*. All resistant lines carried mutations in the apicoplast-encoded large ribosomal subunit RNA (23S rRNA), reminiscent of clindamycin resistance mechanisms found in bacteria. We recovered three different mutations, all located in the peptidyl transferase region of apicoplast 23S rRNA. Each 23S rRNA mutation was associated with >20-fold resistance, although some mutants grew extremely poorly *in vitro* and therefore may lack clinical relevance *in vivo*. We assessed how well our most vigorously growing 23S rRNA mutant could infect *Anopheles* mosquitoes and found a modest reduction in vector infectivity, indicating that high-level clindamycin resistance is likely to be transmissible in the field. This is in contrast to atovaquone resistance, which exhibits a total block to transmission (and hence spread), and azithromycin resistance, which does not significantly impact *P. falciparum* development in the mosquito.

## Introduction

*Plasmodium* resistance to antimalarial treatments undermines malaria control and elimination efforts. The efficacy of excellent antimalarials such as chloroquine, sulfadoxine-pyrimethamine, and artemisinin has been compromised, diminishing the efficacy of these widely-used drugs (1, 2). While promising new antimalarial drug candidates are on the horizon, the spectre of parasite resistance to new drugs remains a concern. Strategies such as drug combinations to combat drug resistance have reduced malaria cases and deaths, but progress has stalled in recent years (3), and multi-drug-resistant parasites are of enormous concern. Innovative and strategic implementation of antimalarial combinations is required to regain control of malaria.

One underexplored strategy for managing antimalarial resistance is to exploit fitness costs associated with mutations that confer parasite drug resistance. Resistance conferring mutations frequently occur in key enzymes or metabolic pathways and may negatively impact parasite growth and transmission (4, 5). Sustained antimalarial use maintains drug-resistant genotypes in the population. However, in the absence of a selection pressure, the evolutionary advantage of drug resistance alleles is lost, and the cost of resistance mutations on fitness negatively impacts the dominance of antimalarial resistant parasites in the population. A classic example is chloroquine resistance in Malawi, where chloroquine resistant parasites all but disappeared following discontinuation of chloroquine use (6). Whilst interesting, reversion to a drug sensitive genotype after cessation of drug use is not a reliable strategy to manage resistance. Instead, we are exploring the fitness costs associated with drug-resistance mutations that prevent or reduce the spread of resistance-conferring alleles via mosquitoes and could, thereby, extend the efficacy of existing antimalarials.

The malaria parasite significantly alters its metabolic activity as it moves between mammalian and insect hosts (7). Mutations conferring resistance to drugs that target pathways with low but essential activity in the blood stages are readily selected by drug pressure, but these mutations can severely impact parasite survival during the mosquito stages, where those pathways become more active (8). This so-called ‘resistance trap’ is evident in mutations in organellar genomes that confer resistance to the antimalarials azithromycin and atovaquone (9–11). Because activity of the apicoplast and mitochondrion increases during the mosquito stages of the parasite life cycle (7, 8), mutations conferring resistance to azithromycin and atovaquone can impair parasite development, thereby limiting transmission of resistance alleles, although the severity of these effects can vary between parasite species (9–11). The impact of this reduction in transmission is further enhanced by maternal inheritance of organellar genomes (12, 13), which prevents complementation or recombination that might otherwise compensate for fitness costs associated with resistant alleles.

Another antimalarial that could fall into this ‘resistance trap’ is the antibiotic clindamycin. Clindamycin, in combination with artemisinin or quinine, is a second line treatment option and, until recently, was used for treatment of malaria in first trimester pregnant women (14). Clindamycin kills malaria parasites by inhibiting apicoplast function (15). It is presumed, based on its mode of action in bacteria (16), that clindamycin binds to the apicoplast-encoded large ribosomal RNA subunit (23S rRNA) and blocks apicoplast protein synthesis (17). As with many other inhibitors of apicoplast housekeeping functions, clindamycin exhibits delayed antimalarial activity (15) and is, therefore, typically administered for malaria treatment as a partner compound alongside fast parasite-killing drugs.

Despite clinical use of clindamycin against malaria, the mechanism(s) and frequency of clindamycin resistance in *Plasmodium* are largely undefined. The only report of clindamycin resistance in *P. falciparum* comes from patient isolates where a mutation in the apicoplast 23S RRNA was postulated to be the mechanism of clindamycin resistance (18). However, whether these resistant isolates were selected by clindamycin treatment for malaria, or arose as bystander resistance during antibiotic treatment for bacterial infections, remains unclear (18). Here, we select *in vitro* for clindamycin resistance in *P. falciparum* to determine if mutations in the apicoplast 23S rRNA are the common mechanism of clindamycin resistance. We investigate the fitness costs associated with resistance mutations in the vertebrate and mosquito hosts to determine the potential for resistant parasites to spread.

## Results and Discussion

### *In vitro* generation of clindamycin resistant *P. falciparum*

To identify the mechanism(s) of clindamycin resistance in *Plasmodium falciparum*, we selected for clindamycin resistance in a multidrug-resistant line (Dd2, Indochina) (19) and a drug-sensitive parasite line (NF54, African) (20, 21). Both strains were independently exposed to either a ‘stepwise’ or a ‘pulsed’ drug selection protocol to identify common mechanisms and mutations that confer clindamycin resistance. We successfully generated four independent clindamycin-resistant lines: *Pf*NF54_Cl^R^_S (stepwise selection), *Pf*Dd2_Cl^R^_S (stepwise selection), *Pf*NF54_Cl^R^_P (pulse selection), and *Pf*Dd2_Cl^R^_P (pulse selection) (Figure 1A). All resistant parasite lines emerged within a few months following the start of selections, indicating that clindamycin resistance is readily attainable *in vitro*.

**Figure 1.**
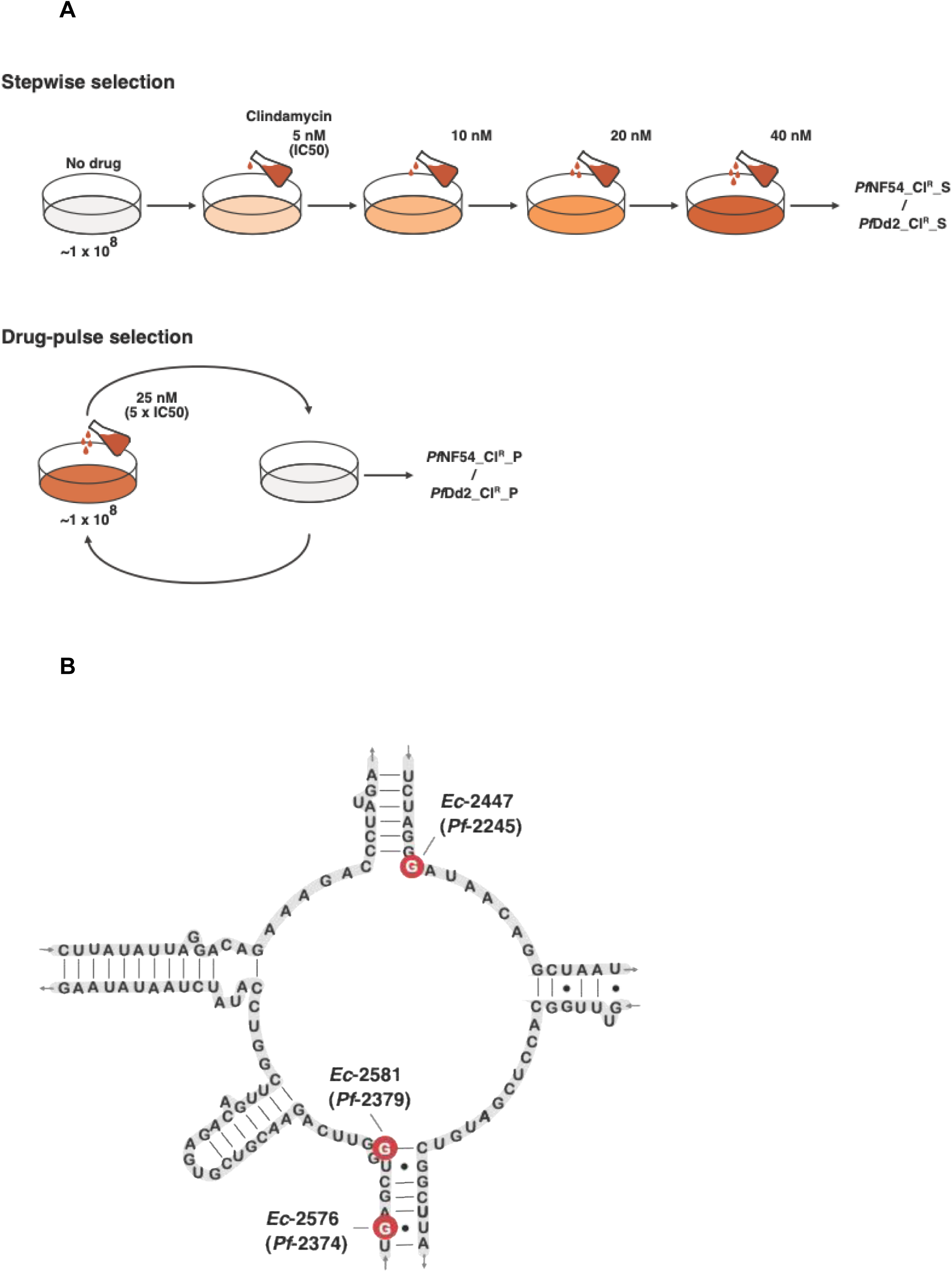
Generation and identification of clindamycin resistant 23S ribosomal RNA *P. falciparum* parasites. A) Graphic of the selection regime methodology to generate *Pf*NF54_Cl^R^_S and *Pf*Dd2_Cl^R^_S by stepwise selection, and *Pf*NF54_Cl^R^_P and *Pf*Dd2_Cl^R^_P by pulse selection. B) Predicted secondary structure of *P. falciparum* peptidyl transferase centre (PTC) of the apicoplast 23S ribosomal RNA as previously published (18). Residues highlighted red indicate location of mutations identified in NF54 and Dd2 selected clindamycin-resistant *P. falciparum*.

### Sequencing analysis

Whole genome sequencing of two clones from each *Pf*NF54_Cl^R^_S and *Pf*NF54_Cl^R^_P independent resistance selections revealed eight mutations across the four clones that were absent from the NF54 drug-sensitive parent line. Seven mutations mapping to pseudogenes were dismissed as being unlikely to contribute to resistance. The remaining mutation was in the apicoplast-encoded 23S rRNA (PF3D7_API04900), a known site of clindamycin resistance in prokaryotes (22, 23). Both clindamycin-resistant lines harbour an identical G → T point mutation at nucleotide position 2245, despite having been selected under different regimens.

Alignment of the *P. falciparum* apicoplast 23S rRNA sequence with homologues shows that G2245 (*Escherichia coli* position 2447) lies in domain V of the large ribosomal subunit (Figure S1), which contains the highly conserved peptidyl transferase centre (PTC), the core catalytic site of the prokaryote ribosome. Note that, for clarity, *E. coli* numbering of the 23SrRNA is used throughout the remainder of this paper unless specified. Modelling of the *P. falciparum* 23S rRNA PTC secondary structure based on *E. coli* situates G2447 within the catalytic core (18) (Figure 1B). Nucleotides that make up the PTC are highly conserved across all domains of life (24) (Figure S1), implying that changes at this region are unusual and potentially significant for antibiotic resistance.

In the apicoplast genome, there are two copies of 23S rRNA located in inverted repeats (25). Whole genome sequencing of resistant lines indicated that both copies of 23S rRNA contain the G2447T mutation. To confirm this, we performed amplicon sequencing of inverted repeat A and inverted repeat B independently (Figure S2).

Both *Pf*NF54_Cl^R^_S and *Pf*NF54_Cl^R^_P contain the G2447T mutation in both their apicoplast 23S rRNA genes (Figure S2). An inverted repeat containing rRNA genes is a canonical feature of plastid genomes from plants and protists, including malaria parasites. Identity of the plastid inverted repeats is maintained by gene conversion (26). We postulate that an initial clindamycin resistance-conferring mutation arose in one apicoplast rRNA gene and was subsequently propagated into the other repeat by gene conversion under drug selection pressure.

Amplicon sequencing of the apicoplast 23S rRNA from *Pf*Dd2_Cl^R^_S and *Pf*Dd2_Cl^R^_P also revealed point mutations in the PTC region of the apicoplast 23S rRNA. *Pf*Dd2_Cl^R^_S contain a G → T mutation at position 2581 (*Pf*-2379).

*Pf*Dd2_Cl^R^_P harbours a G → T mutation five nucleotides upstream at position 2576 (*Pf*-2374). Like G2447T (the mutation recovered twice after *Pf*NF54 selection, see above), G2581 and G2576 are also conserved across *E. coli*, the plastid in the photosynthetic apicomplexan relative *Chromera velia,* and in other apicoplast-containing apicomplexan species (Figure S1). Extremely slow *in vitro* asexual growth of *Pf*Dd2_Cl^R^_P prevented generation of clonal lines required for whole genome sequencing. Therefore, we cannot exclude the possibility of other mutations outside the 23S rRNA that confer, or enhance, clindamycin resistance.

Unfortunately, the apicoplast genome cannot be genetically modified, so recapitulation of the putative resistance mutations in a clean genetic background is not possible. Nevertheless, given the absence of biologically relevant shared mutations outside the apicoplast 23sRNA in the fully sequenced NF54 lines, the most parsimonious explanation is that mutations in the apicoplast-encoded 23S rRNA in all our selected parasite lines are the primary source of clindamycin resistance.

### *Pf*NF54_Cl^R^_S/P and *Pf*Dd2_Cl^R^_S/P are highly resistant to clindamycin but not to other inhibitors of apicoplast translation

To determine the degree of clindamycin resistance in *Pf*NF54_Cl^R^_S/P and *Pf*Dd2_Cl^R^_S/P, we quantified clindamycin IC50 values for our mutant lines for comparison with their parental lines. All four independent clindamycin resistant parasite lines exhibited greater than 20-fold increased resistance to clindamycin (Figure 2A, B). *Pf*NF54_Cl^R^_S and *Pf*NF54_Cl^R^_P showed similar mean IC50s of 170 nM and 182 nM respectively, which is consistent with them harbouring the same G2247T 23S rRNA mutation. *Pf*Dd2_Cl^R^_S and *Pf*Dd2_Cl^R^_P had mean IC50s of 154 nM and 207 nM respectively. As mentioned, despite multiple attempts, we were unable to clone *Pf*Dd2_Cl^R^_P parasites due to significant growth defects. Therefore, drug assays and subsequent phenotyping for this line were performed on non-clonal populations. The difference in IC50 values of *Pf*Dd2_Cl^R^_S and *Pf*Dd2_Cl^R^_P is likely due to different 23S rRNA mutations (Fig. 1B), but we cannot rule out the possibility that additional resistance-conferring or compensatory mutations exist elsewhere in the genome for *Pf*Dd2_Cl^R^_S and *Pf*Dd2_Cl^R^_P lines.

**Figure 2.**
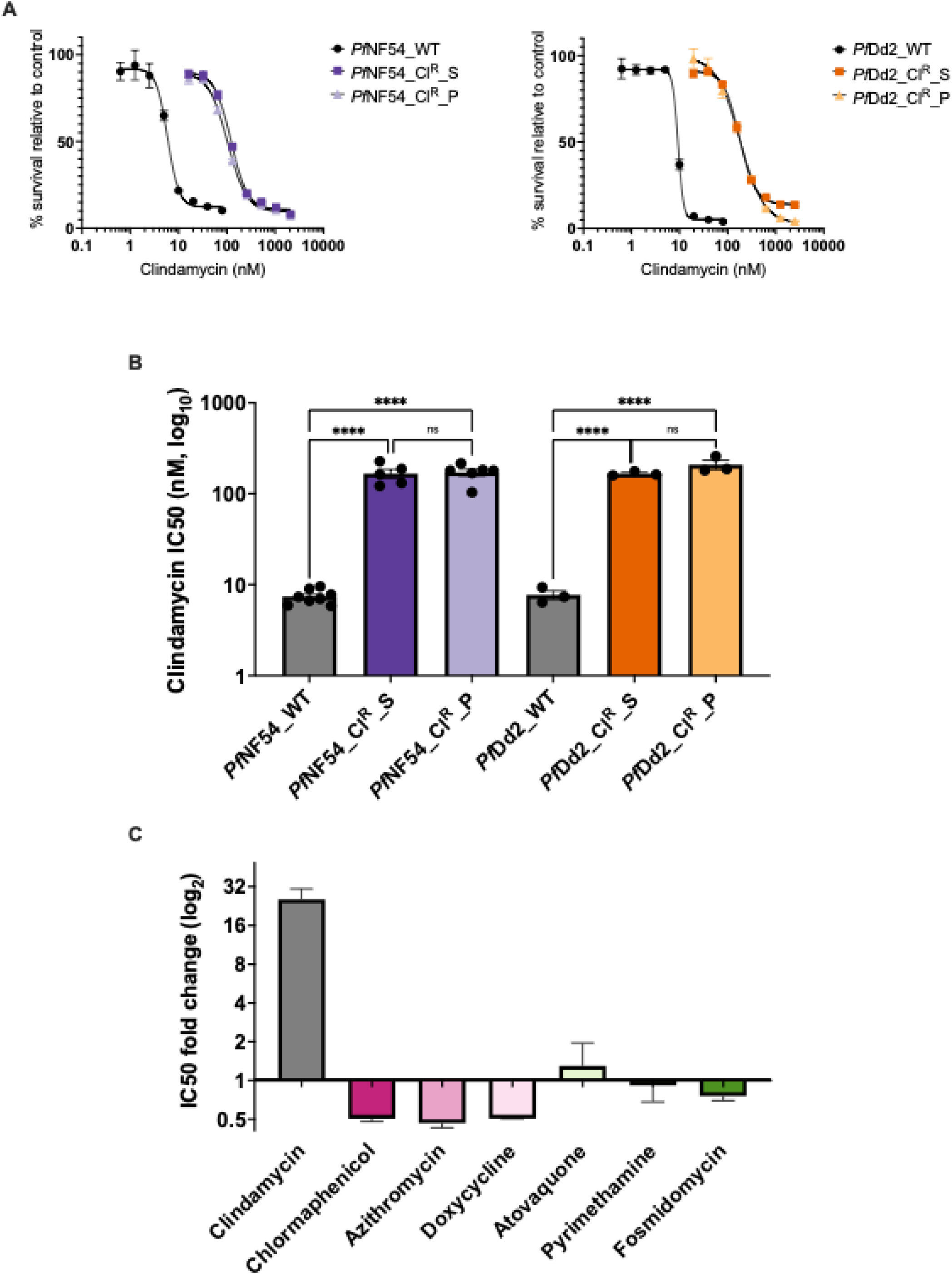
Asexual and sexual growth of *Pf*NF54 and *Pf*Dd2 clindamycin resistant parasites. A) Representative clindamycin dose response curves of *Pf*NF54_Cl^R^_S/P and *Pf*Dd2_Cl^R^_S/P relative to a no-drug control. *Pf*NF54 clindamycin drug assays were performed over 144 hours whilst *Pf*Dd2 assays were performed over 168 hours. Error bars represent the standard deviation. B) Comparison of growth inhibition (IC50) among NF54 parasite lines (*Pf*NF54_Cl^R^_S (n=5), *Pf*NF54_Cl^R^_P (n=6) and *Pf*NF54_WT (n=8)) and Dd2 parasite lines (*Pf*Dd2_Cl^R^_S (n=3), *Pf*Dd2_Cl^R^_P (n=3) and *Pf*Dd2_WT (n=3)) across ‘n’ biological replicates. Statistical significance was determined by one-way ANOVA (****=P≤0.0001). Error bars represent the mean ±SEM. C) IC50 fold change of *Pf*NF54_Cl^R^_S from control parasites to apicoplast and non-apicoplast translational inhibitors from five independent experiments for clindamycin and two independent experiments for remaining antimalarials. Error bars represent the geometric mean with geometric SD.

Mutations at residues G2447 (27–32), G2576 (32, 33) and G2581 (32) of 23 rRNAs also confer resistance to chloramphenicol, linezolid and anisomycin in select species of Bacteria and Archaea. Moreover, mutations or modifications at the binding site of clindamycin in the 23S rRNA can confer high level cross-resistance to other translation-targeting antibiotics (34). Therefore, we assayed our selected clindamycin resistant lines against chloramphenicol and other putative apicoplast translational inhibitors to test for cross-resistance. Linezolid and anisomycin were not tested in this study as linezolid is not active against *P. falciparum* (35), and anisomycin is a universal rRNA inhibitor targeting both the cytoplasmic and organellar translational machineries of *Plasmodium* (36).

*Pf*NF54_Cl^R^_S and *Pf*NF54_Cl^R^_P exhibited no increased resistance against any of the tested antimalarials other than clindamycin. Intriguingly, both NF54 clindamycin resistant lines were approximately 2-fold more sensitive to three antibiotics that are known or predicted to target apicoplast translation: namely, azithromycin, doxycycline, and chloramphenicol (Figure 2C, Table S1)(15, 17, 37). A similar, slight increase in antibiotic sensitivity was reported in azithromycin resistant 7G8 *P. falciparum* parasites when tested against tetracycline, doxycycline, and thiostrepton compared to the parental line (38). The authors hypothesise that point mutations in 23S rRNA could change ribosomal structure and allow antibiotics with distant binding sites on the ribosome to be slightly more efficacious (38). Another potential reason for the observed increased sensitivity to other antibiotics in clindamycin-resistant mutants is that 23S rRNA mutations could alter apicoplast translation rates, thus rendering parasites more sensitive to other apicoplast translation-targeting antimalarials. Although the cause of hypersensitivity to apicoplast translational inhibitors is unclear, this observation could make clindamycin and azithromycin useful partner drugs or candidates for mass drug administration if the reported mechanisms of resistance are mirrored in natural populations.

### Asexual growth of clindamycin resistant Dd2 parasites is slow

Resistance conferring mutations often arise at a cost to parasite fitness (4). In a drug-free environment, resistant parasites can be at an evolutionary disadvantage when competing against wild type (drug-sensitive parasites), which in turn can reduce their likelihood for transmission and spread. To assess the fitness costs associated with clindamycin resistance, we monitored the asexual growth over four intraerythrocytic developmental cycles of NF54 and Dd2 clindamycin resistant parasites in conjunction with their respective parental lines using a SYBR Green 1-based fluorescence assay (39). The asexual replication of *Pf*Dd2_Cl^R^_S and *Pf*Dd2_Cl^R^_P is markedly reduced compared to the parental control, whilst both NF54 clindamycin resistant parasites retain wild type rates of asexual growth (Figure 3A). These observations align with the impact of corresponding mutations on growth in Bacteria and species of Archaea. G2447 mutants have very mild to no growth defects (27, 28, 31, 40, 41), whilst growth of G2576 and G2581 mutants is significantly reduced compared to wild type in *Mycobacterium smegmatis, Streptococcus pneumoniae,* and *E. coli* (33, 42, 43).

**Figure 3.**
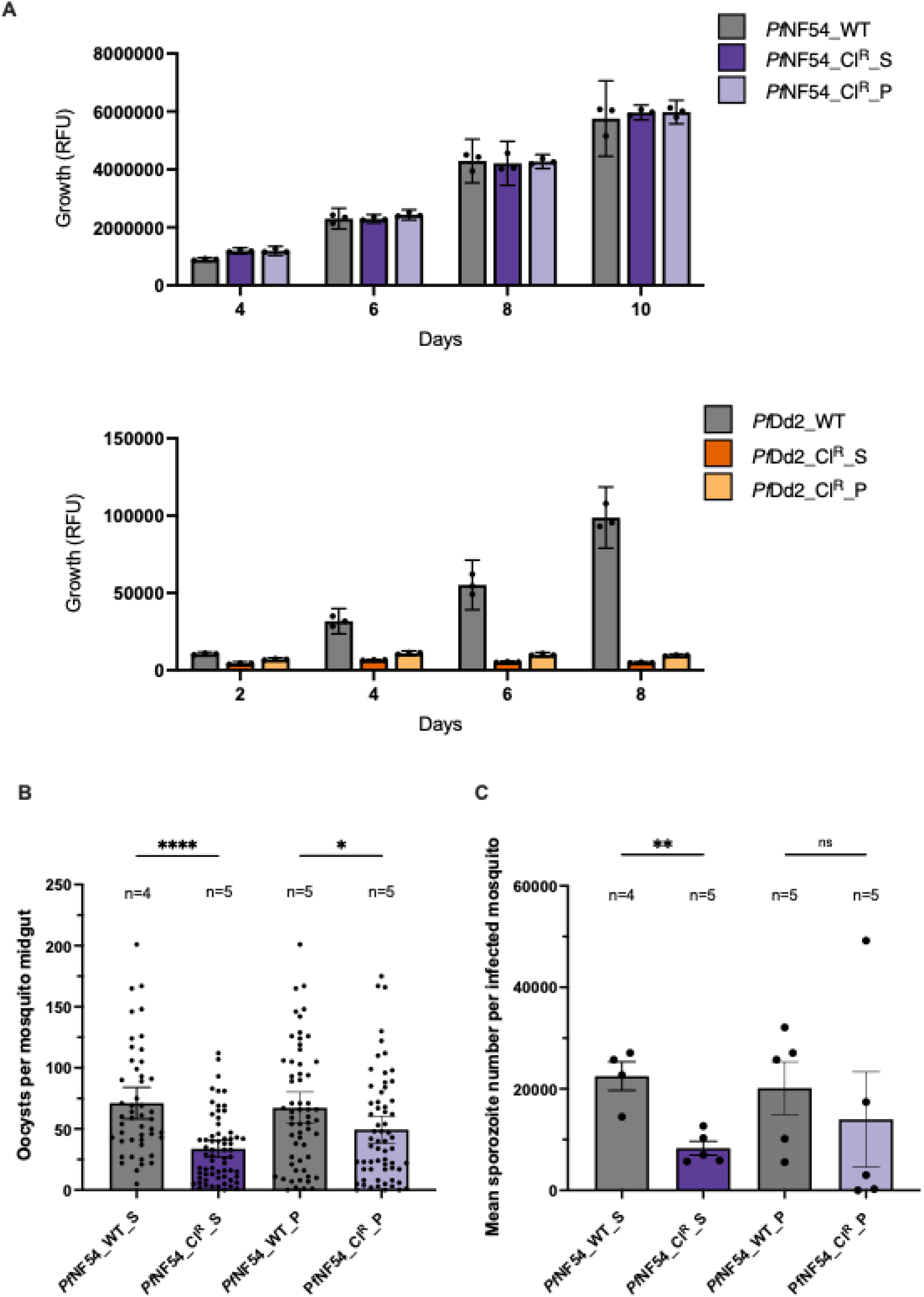
Asexual and sexual growth of NF54 and Dd2 clindamycin resistant parasites. A) Asexual growth of clindamycin resistant NF54 and Dd2 parasite lines. Ring-stage parasites were plated in triplicate at a starting parasitemia of 0.5% and split 1:2 (*Pf*NF54) or 1:4 (*Pf*Dd2) every 48 hours. Blood stage growth, measured by relative fluorescence units (RFU), of clindamycin resistant parasites was determined relative to the wildtype control across four cycles using a SYBR Green I-based fluorescence assay. Error bars represent the mean ±95% confidence interval. B) *Pf*NF54_Cl^R^_S and *Pf*NF54_Cl^R^_P are impacted in the development of mosquito midgut oocysts. Control infections for *Pf*NF54_Cl^R^_S and *Pf*NF54_Cl^R^_P are shown as *Pf*NF54_WT_S and *Pf*NF54_WT_P respectively. A minimum of 10 midguts from *A. stephensi* mosquitoes infected with *Pf*NF54_WT_S, *Pf*NF54_Cl^R^_S, *Pf*NF54_WT_P and *Pf*NF54_Cl^R^_P or *Pf*NF54_Cl^R^_P were dissected per independent infection (‘n’) at either day 7- or 8-post mosquito blood feed for oocyst counts. Statistical significance was determined using the Mann-Whitney non-parametric test (****=P≤0.0001, *=P≤0.05) and error bars represent the median ±95% confidence interval. C) *Pf*NF54_Cl^R^_S, but not *Pf*NF54_Cl^R^_P, produce significantly fewer salivary gland sporozoites compared to control parasites. *Pf*NF54_WT_S and *Pf*NF54_WT_P represent control infections for *Pf*NF54_Cl^R^_S and *Pf*NF54_Cl^R^_P respectively. The mean sporozoite number per independent experiment (‘n’) was obtained by dissecting salivary glands of at least 10 infected mosquitoes. Statistical significance was determined using paired t-test (**=P≤0.01), error bars represent the mean ±SEM.

Considering the seemingly normal asexual development of NF54 clindamycin resistant parasites, it is surprising that the G2447 mutation was not readily selected for in Dd2 parasites. This anomaly could be random variability or might suggest that the fitness costs associated with particular residue changes in the 23S rRNA differ between the NF54 and Dd2 parasite strains. Whatever the case, it does highlight the effect of varying parasite genetic backgrounds when pursuing *in vitro* evolutionary studies of antimalarial resistance. The stark growth defect of *Pf*Dd2_Cl^R^_S and *Pf*Dd2_Cl^R^_P suggests that clindamycin resistance associated with mutations at 23S rRNA G2581 or G2576 are unlikely to compete in wild populations, so may not be clinically relevant.

### Clindamycin resistant NF54 parasites have development defects in mosquitoes

To assess the ability for clindamycin resistance to transmit, we tested the sexual viability of our selected NF54 clindamycin resistant mutants, which exhibit robust resistance and asexual growth. The mosquito infectivity of culture-adapted Dd2 parasites is poor (44), and therefore *Pf*Dd2_Cl^R^_S and *Pf*Dd2_Cl^R^_P were only analysed in the asexual blood stages. However, the impact of G2581 and G2576 23S rRNA mutations on asexual replication likely limits subsequent transmission of resistant *P. falciparum* in wild populations anyway.

To determine whether NF54 clindamycin resistant parasites can establish an infection in the mosquito host and subsequently produce infectious sporozoites, we allowed our cultures to develop gametocytes and fed the blood to *Anopheles stephensi* mosquitoes. Mosquito midguts were examined at approximately day 7 post blood feed, and mosquito salivary glands at day 18, to quantify the numbers of *Plasmodium* parasites per mosquito.

Mosquitoes infected with *Pf*NF54_Cl^R^_S produced significantly fewer midgut oocysts and salivary gland sporozoites compared to mosquitoes infected with the parental (clindamycin sensitive) parasite line (Figure 3B, C). This reduction was consistent among both clindamycin resistant lines however, variability in the salivary gland sporozoite numbers of *Pf*NF54_Cl^R^_P precluded conclusions about statistical significance. Whereas atovaquone-resistant *P. falciparum* produce zero sporozoites in mosquitoes (10, 11), our clindamycin-resistant produce fewer sporozoites compared to sensitive, wild type lines. Thus, G2447T 23S rRNA mutants are potentially able to transmit but perhaps with reduced efficacy compared to drug sensitive parasite populations in the wild.

### Concluding remarks

Clindamycin has a good safety profile and is useful in treating malaria infections in vulnerable populations (45–48), and combination-based therapies including clindamycin are effective (45, 47). Our exploration of malaria parasite clindamycin resistance substantially extends previous work (18) and flags mutations in the peptidyl transferase centre of the apicoplast 23S rRNA as a ‘hotspot’ for mutations associated with clindamycin resistance. Although the G2447T mutation in the 23S rRNA is the only biologically relevant mutation shared between our fully sequenced resistant parasites, we cannot experimentally confirm that 23S rRNA mutations alone are sufficient to confer clindamycin resistance because genes in the apicoplast cannot currently be edited. Nevertheless, the four resistant lines identified here, plus another mutation published previously (18), all have 23S rRNA mutations and no other mechanism of clindamycin resistance in malaria parasites is known. Thus, any clindamycin resistance surveillance strategy should include apicoplast 23S rRNA sequencing as a minimum.

Our NF54 clindamycin resistant parasites, with mutations at G2447, grew at wild type rates in the blood stage. In contrast, clindamycin resistant Dd2 parasites that harbour different 23S rRNA mutations were markedly slower growing. We speculate that the mutations found in the Dd2 parasites may not be stable *in vivo*, where slow parasite growth favours the host in eliminating infection. Why we recovered different sets of 23S rRNA from different strains undergoing the same selection regimes is not known, but it does highlight the usefulness of not relying on a single strain to explore resistance mechanisms.

Mosquito infectivity assays of parasites harbouring the 23S rRNA mutation associated with clindamycin resistance indicate a modest fitness loss in the insect host. We conclude that this mild reduction in parasite mosquito stage fitness would likely still allow the 23S rRNA mutation to be transmitted and hence spread, especially considering that a relatively small inoculum of sporozoites is sufficient to achieve infection of vertebrate hosts (49). However, further investigation into the impact of the 23S rRNA mutation on parasite liver stage development could identify additional impairments to transmission competency, particularly since *P. falciparum* azithromycin resistant mutants exhibit a defective exoerythrocytic development in humanised mice (9). It remains unknown whether the A2058C mutation found in clindamycin resistant *P. falciparum* isolates (18) will transmit efficiently via mosquitoes.

We commenced this work to explore the fitness costs imposed by clindamycin resistance and how that impacts the ability of drug-resistant human malaria parasites to survive and spread. We found several 23S rRNA mutations associated with clindamycin resistance but only one (G2447T) that allows for robust blood stage growth. The G2447T mutants exhibit a modest inhibition of mosquito infectivity and is likely to transmit. This contrasts with the absolute transmission block seen in atovaquone-resistant parasite lines (10, 11), but is a more pronounced effect than azithromycin-resistant *P. falciparum*, which suffers no diminution of parasite numbers in mosquitoes (9). Transmission studies thus far indicate that resistance to apicoplast targeting drugs (clindamycin and azithromycin) is more likely to spread than resistance to the mitochondrial inhibitor atovaquone, which have a total transmission block. Assaying fitness of drug-resistant mutants is thus essential to estimate their ability to spread and judge whether they will persist in natural populations in the absence of drug pressure.

## Materials and Methods

### Drugs

Clindamycin hydrochloride monohydrate (CAS Number: 58207-19-5), Azithromycin dihydrate (CAS Number: 117772-70-0), doxycycline hyclate (CAS Number: 24390-14-5) and atovaquone (CAS Number: 95233-18-4) were acquired from Sigma-Aldrich. Pyrimethamine was obtained from Tokyo Chemical Industry Co. (CAS Number: 58-14-0) and fosmidomycin was kindly provided to us by Stuart Ralph. Antimalarial drugs were diluted in either dH_2_O, ethanol or DMSO according to compound solubility to make 1 mM stocks stored at -20 °C. Working stocks were further diluted in RPMI 1640 for experiments as required.

### Parasite culturing

NF54 and Dd2 parasites were cultured at 4% haematocrit in RPMI 1640 supplemented with 5% w/v Albumax, 25 mg/mL hypoxanthine in 1M NaOH and 10 mg/mL gentamycin. Red blood cells (type O) and human serum are kindly donated by the Australian Red Cross Lifeblood. Parasites were maintained as described previously (50) and stored at 37 °C in perspex boxes gassed with 5% CO_2_ + 1% O_2_ + balance N_2_ (BOC).

### Resistance selection

Clindamycin resistance was selected for by treating a clonal population of approximately 1 x 10^8^ *P. falciparum* Dd2 or NF54 parasites to clindamycin. In stepwise selections, parasites were exposed to 5 nM clindamycin and drug pressure was applied continuously throughout the resistance selection. The concentration of clindamycin was doubled when parasites became accustomed to the previous dose. Drug pulse selections were performed by exposing parasites to 5x IC50 concentration of clindamycin (25 ng). Clindamycin was removed from the culture when the parasitemia began to decline, allowing cultures to recover in drug-free media. Clindamycin (25 ng) was reintroduced to the culture when the parasitemia reestablished. This cycle was continued until parasites grew stably in the presence of 25 ng clindamycin or above. All parasite lines were cloned by limiting dilution except *Pf*Dd2_CLINDR_2 and resistance was confirmed by *in vitro* SYBR Green I drug assays.

### *In vitro* drug assays

Parasite susceptibility to antimalarial drugs were performed in a 96-well plate and parasites survival was measured using the SYBRGreen I fluorescence based system (39). Synchronised ring-stage parasites were plated in triplicate (0.025% parasitemia for *Pf*Dd2_CLINDR_1 and *Pf*Dd2_CLINDR_2, 0.05% parasitemia for clindamycin, azithromycin and doxycycline, or 0.125% parasitemia for atovaquone, fosmidomycin and pyrimethamine drug trials) at 2% haematocrit to serially diluted antimalarial compounds. Parasites were exposed to drug for 72 hours in atovaquone, fosmidomycin and pyrimethamine drug trials, 144 hours in clindamycin, azithromycin and doxycycline drug trials, or 168 hours in trials testing drug susceptibility in *Pf*Dd2_CLINDR_1 and *Pf*Dd2_CLINDR_2 specifically due to slow asexual growth of these parasite lines. In the 144 and 168-hour assays, culture media was replaced with drug-free media at 72 hours and 120 hours. At the end of the assay, cells were incubated in lysis buffer containing SYBR green (0.0001%) for 1 hour and read on the CLARIOstar Plus (BMG LABTECH) plate reader to determine parasite survival relative to the drug-free control parasites.

### *In vitro* growth assays

Highly synchronous ring-stage parasites were plated in triplicate at 0.125% parasitemia and 2% hematocrit in a 96-well plate. Every 48 hours, 100 μL of parasite culture was transferred to a new 96-well plate for growth analysis. The remaining culture was refreshed with 2% hematocrit to result in a 2:1 dilution for NF54 parasite lines or subbed in half again before refreshing with 2% hematocrit to result in a 4:1 dilution for Dd2 parasite lines. Multicycle growth analysis was performed by lysing cells with buffer containing SYBR green (0.0001%) and incubating for 1 hour before measuring the relative fluorescence on the CLARIOstar Plus (BMG LABTECH) plate reader.

### Whole genome sequencing

DNA from two clonal *Pf*nf54_CLINDR_1 and *Pf*nf54_CLINDR_1, and the parental *Pf*nf54_WT line, was prepared for whole genome sequencing using the QIAamp DNA Blood Mini Kit (QIAGEN) following manufacturer’s instructions. Genomic DNA samples were sent to the VCGS (Murdoch Children’s Research Institute, Royal Children’s Hospital, Parkville, Victoria) for Illumina DNA library preparation and sequenced at 8 M reads (∼100x coverage) per sample on the NovaSeq6000 (2x150 bp reads). Reads were mapped using the Burrows-Wheeler Aligner (51) to the *P. falciparum* 3D7 genome version 68, obtained from PlasmoDB (52). Mapped reads were visualised in IGV (53), and checked with FastQC (54) and Qualimap (55). Mutations were identified using GATK following the best-practices documentation (56) then classified using SnpEff and SnpSift (57, 58). Further analysis was performed using GNU/Linux coreutils and the AWK programming language (59). Copy number variations were detected using FREEC (60), with further analysis using GNU/Linux coreutils.

### Gametocyte culturing and mosquito infections

Infectious stage V *P. falciparum* NF54 gametocytes were cultured in six-well plates at 5% haematocrit in RPMI 1640 media supplemented with 5% human serum and 25 mg/mL hypoxanthine in 1M NaOH. Cultures were set up at a starting asexual parasitemia of 0.5% and media was changed slowly, without the addition of fresh blood, daily for 18 days. *Anopheles stephensi* mosquitoes, which were reared and maintained as previously described (61), were infected with 0.15% stage V gametocytes using glass membrane feeders. Mosquitoes were starved for 48 hours post blood-feed to kill any mosquitoes that did not feed, before providing cotton wall soaked in a 10% sucrose solution to the mosquito cage.

### Mosquito dissections

Midguts of infected *A. stephensi* mosquitoes were dissected at day 7-8 post blood feed and stained with 0.2% mercurochrome for 10 minutes followed by a wash with 1x dPBS for 10 minutes before mounting on to a slide. The number of oocysts per mosquito midgut was determined. Salivary glands of blood-fed mosquitoes were dissected on day 17-19 post blood feed for salivary gland sporozoites. The average number of sporozoites of a pool of 10-20 mosquitoes was counted on a Neubauer haemocytometer. The midguts of mosquitoes dissected for salivary gland sporozoites was also dissected to confirm mosquito was infected and determine the average number of sporozoites per infected mosquito.

### Statistical analysis

All phenotypic data were analysed using Prism 10 software (GraphPad Prism version 10.6.1). IC50 concentrations were determined using the variable slope (four parameters) equation and statistical significance of IC50s between parasite lines was determined by one-way ordinary ANOVA. Significance between drug sensitive and clindamycin resistant mosquito oocysts and sporozoites was determined using Mann-Whitney non-parametric test and paired t-test respectively. Results were considered significant when P<0.05.

